# Viral RNAs as Dual Graphs: Extending the Motif Universe of RNAs

**DOI:** 10.64898/2026.06.20.733508

**Authors:** Jad Zbib, Tamar Schlick

**Author notes:** Corresponding author: Tamar Schlick.

## Abstract

In the evolving landscape of RNA research, the classification and analysis of RNA motifs is necessary to uncover the intricate mechanisms governing cellular and viral processes. Here we apply the coarse-grained RAG (RNA-As-Graphs) framework to advance the classification and understanding of RNA motifs, with a focus on expanding the RNA Motif Atlas through the inclusion of novel viral RNA structures. By analyzing 273 experimentally resolved viral RNA structures from the Protein Data Bank (PDB) using RAG dual-graph representations, we identify 14 previously uncatalogued viral RNA motifs. These motifs, which include tRNA mimicking domains, exoribonuclease-resistant domains, and internal ribosome entry sites, expand the diversity of RNA to a total number of 197 dual graph motifs. We applied k-means, PAM, and Ward clustering and observed substantial overlap between viral and general RNAs. The expanded library of RNA motifs and submotifs provides a resource for motif discovery and RNA design.

## 1 Introduction

In recent years, RNA has enjoyed heightened focus from the scientific community, as evidenced by the expanding research in RNA structure, function, and interactions. Extensive research has been dedicated to unraveling the complexities of coding and noncoding RNA, including interactions with DNA and proteins, and their role in gene expression and genome maintenance (1). The intricate involvement of different types of RNA in numerous cellular and biological processes has also drawn attention to RNAs as potential therapeutic targets. Notable examples are the mRNA-based vaccines for COVID-19, such as those developed by Pfizer and Moderna. In addition, a range of siRNA, miRNA, and aptamer RNA based therapeutics are in clinical development, with some granted FDA approval (2; 3). RNA therapies have the advantage of having no significant genotoxic effects as compared to DNA therapies (4), and could define safer direct genome based interventions.

RNA secondary structure refers to paired and unpaired ribonucleotides as folded in the single-stranded polynucleotide chain. These nucleotide pairings are driven by various canonical and non-canonical types of base-base interactions, such as Watson-Crick, Hoogsteen, or sugar edge (5). Each type of pairing may present stereoisomeric variations (cis and trans pairing). RNA secondary structure often contains defined motifs, such as hairpin loops, pseudoknots, and quadruplexes, and such motifs are employed in many biological processes. A shared structural basis can be associated with different functions, while similar biological functions, such as transcription initiation and nuclease resistance, can also exploit a variety of RNA motifs (6; 7). This versatility underscores the complexity and variability in RNA behavior, making RNA an interesting subject for computational biology research.

Coarse-grained models, such as graph representations, are useful for representing RNAs due to their ability to simplify complex molecules while retaining key structural and functional characteristics of interest. Such modelsuce computational complexity by grouping smaller building blocks of molecules into larger units, thus reducing the number of degrees of freedom and accelerating the study of larger RNA molecules. In the 1970s and 1980s, pioneers like Waterman, Shapiro, Nussinov, and others introduced various representations where RNA elements such as stems, single strands, and loops were assigned to different graph objects. These early works used graphs to model and define RNA secondary structures, representing multiple base pairs as edges and single-stranded RNA as vertices (8; 9).

These representations facilitated structural comparisons, and graph derived objects invited novel mathematical approaches to analyze and describe RNA structure, such as approximating the free energy of RNA structures using adjacency matrices (8; 10). Early graph representations fell short in capturing complex motifs, most notably pseudoknots, which are critical in fully understanding RNA’s biological functions. In 2003 (11), dual graphs for representing RNA were introduced by the Schlick group, enabling the representation of pseudoknots. By representing RNA structures as graphs, where nodes denote stems, and edges denote single-stranded RNA, we can capture the essential topological and geometric features of RNA. This abstraction facilitates the analysis of RNA folding, dynamics, assembly and interactions, making it easier to identify patterns and motifs critical for biological functions, as exploited recently to estimate the size of the RNA motif universe (21). Such reduced models turned out especially useful for designing minimal mutations for the SARS-CoV-2 frameshifting element and describing folding landscapes (19; 20; 43).

The development of the RNA-as-Graphs (RAG) Motif Atlas (11) began with the more intuitive tree graphs, where loops are vertices and stems are connecting edges. However, tree graphs cannot represent pseudoknots. A pseudoknot is a secondary structure 3D motif formed when a single stranded loop in a hairpin forms base pairs with another single stranded region, possibly another distal hairpin loop. Pseudoknots are recurring motifs in both coding and non-coding RNA, and they play roles varying from regulation of translation initiation to controlling translation elongation and frameshifting (32; 35). To represent pseudoknots, dual graphs employ the opposite convention where loops are edges and stems are vertices. Figure 1 shows the dual graph of a three-way junction RNA and Figure 2 shows the graph of a large more complex pseudoknotted RNA. Using dual graphs, pseudoknots (loop-loop or loop-stem bonding between two hairpin structures) can be represented, as shown in Figure 2.

**Figure 1:**
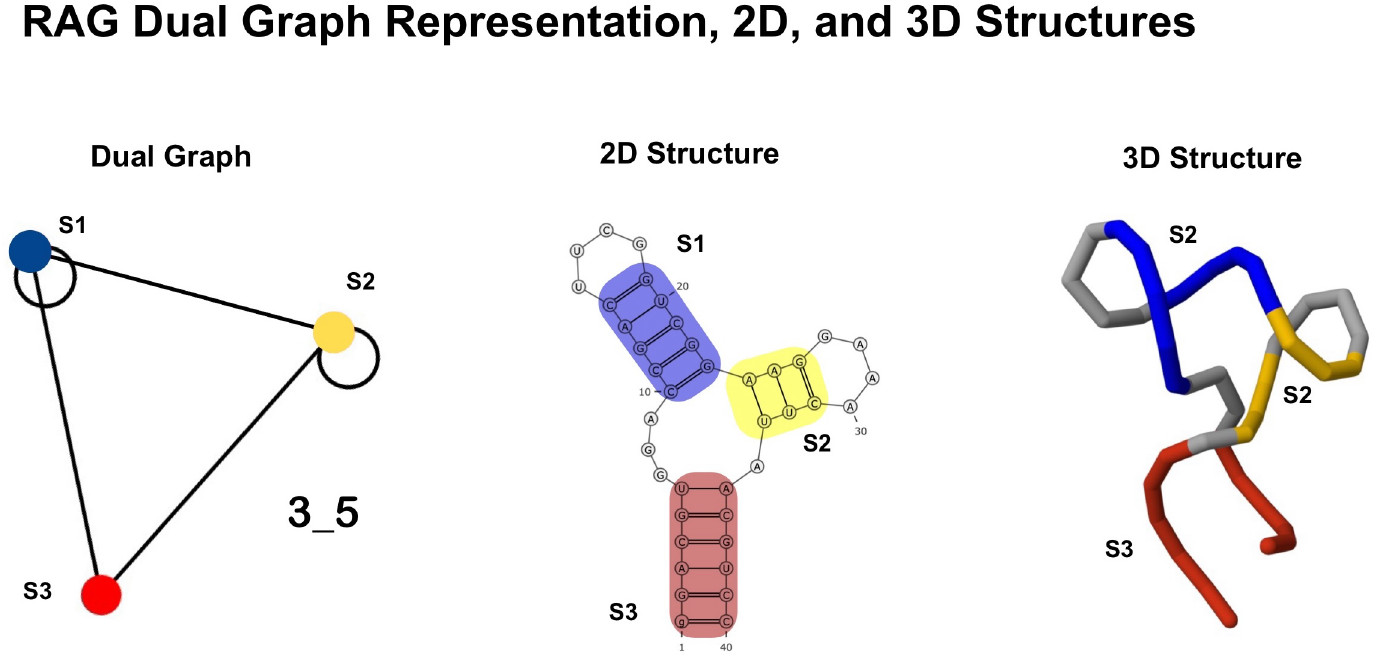
A selected three-way junction-containing RNA from dengue fever virus RNA (PDB ID: 7UMD), shown in 3D, 2D, and dual-graph representation (Graph ID 3_5). The three stems are colored and labeled across all panels (S1 yellow, S2 blue, S3 red), and gray regions denote loops, matching each stem in the RNA substructure to its corresponding vertex in the dual graph.

**Figure 2:**
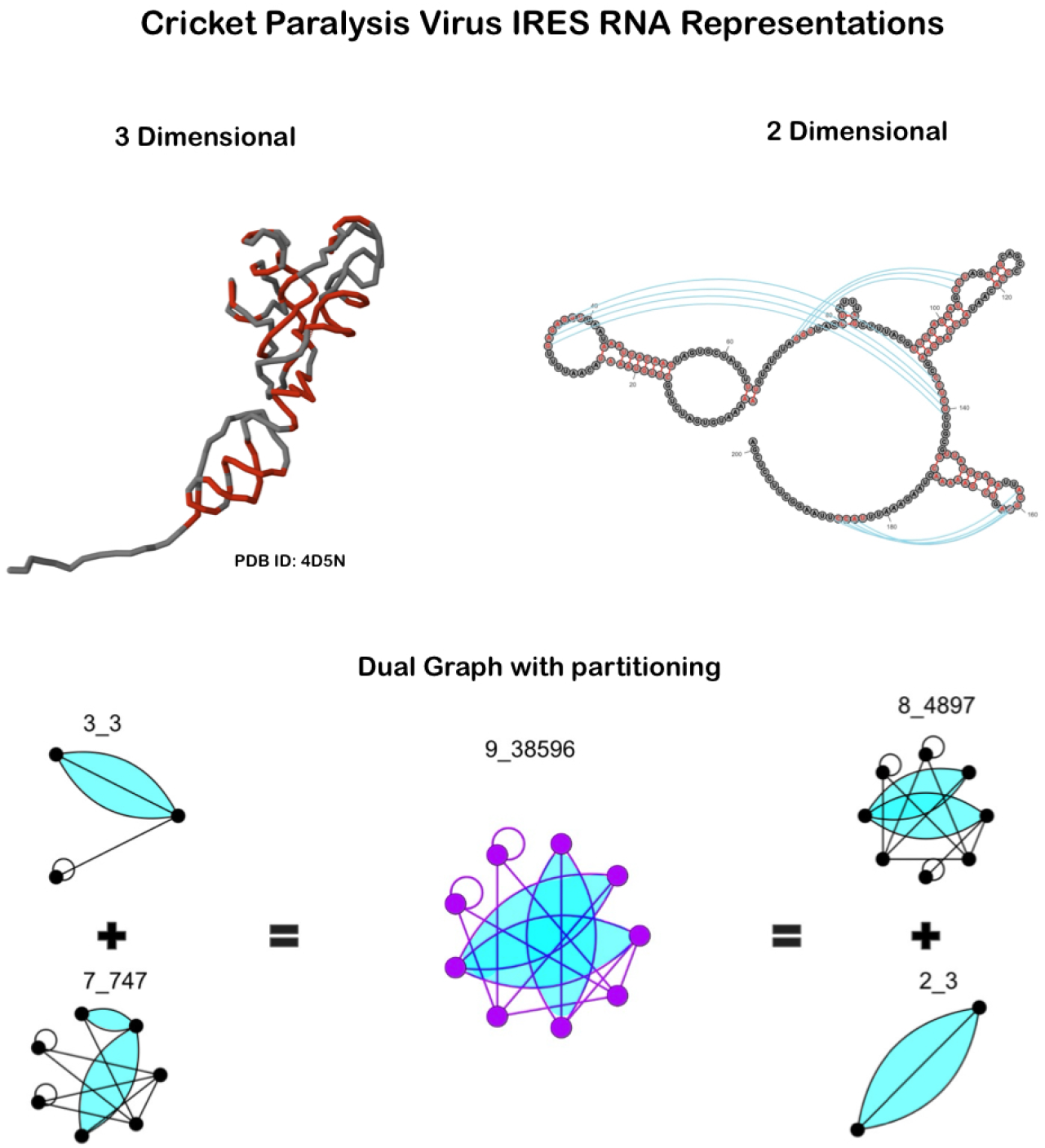
3D, 2D, and dual graph 9_38596 representations of the Cricket Paralysis Virus IRES (PDB ID: 4D5N) and its partitioning. In the 3D and 2D structures, red nucleotides indicate base pairs forming a stem. Two different partitionings of dual graph 9_38596 are shown, with intact pseudoknots highlighted in light blue. The first partitioning results in graphs 7_747 and 3_3, while the second leads to graphs 8_4897 and 2_3.

While RNA generally serves as a versatile agent for carrying genetic information and participating in various cellular processes, viral RNAs likely possess unique characteristics due to distinct strategies for maintaining and propagating a viral life cycle.

Here we expand the RAG motif atlas, adding 14 previously uncatalogued viral motifs derived from 273 viral associated RNA and increase the RNA motif atlas from 183 to 197 existing motifs (24). Next, to assess whether viral RNAs occupy a distinct region, we cluster all RNA motifs using k-means (25), PAM (26), and Ward (27) (See Methods). The resulting clustering show that viral and general RNAs do not occupy distinct clusters in RNA motif space and that they overlap, indicating a sharing of basic RNA submotifs. Finally, we highlight the shared motifs between viral and general RNAs, which rely on common secondary structure building blocks.

## 2 Results

### 2.1 Viral Graph Motifs and Submotifs

Our expanded motif atlas containing viral RNAs in Figure 3 was formed for the addition of viral RNAs. We identified 273 distinct PDB files of viral RNA structures using the Research Collaboratory for Structural Bioinformatics (RCSB) (31) MeSH search. The PDB entries had entity types RNA AND correspond to a natural source organism (to exclude synthetic structures) AND they also belong to a viral taxonomy represented in the PDB, including Riboviria, Duplodnaviria, Varidnaviria, Ribozyviria, unclassified bacterial viruses, and unclassified sequences.

**Figure 3:**
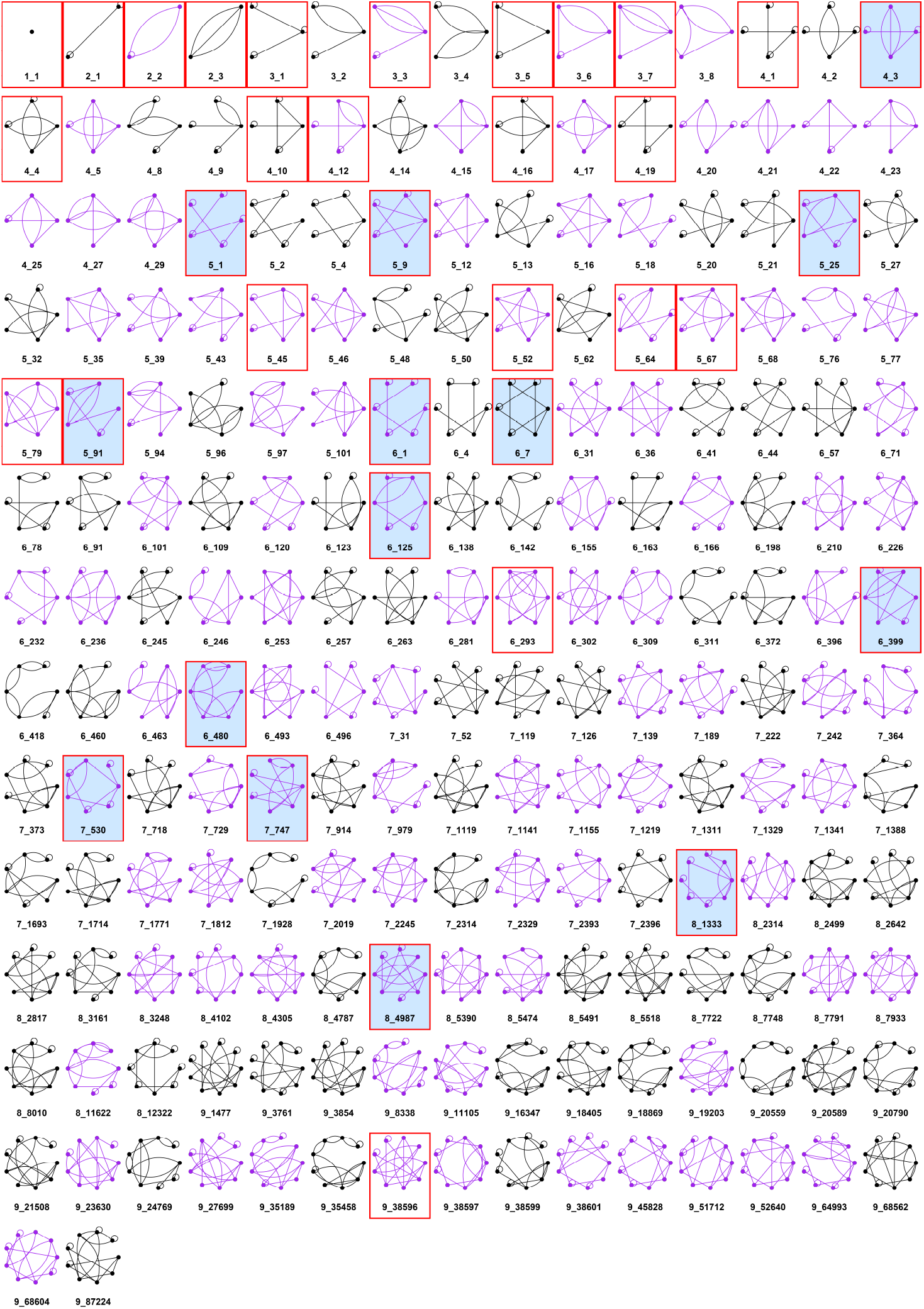
The assembled library of 197 dual graphs in the updated RAG Atlas, comprising viral and general RNAs. Graphs containing at least one pseudoknot are highlighted in purple. All 37 distinct viral dual graphs identified in our dataset are boxed in red, while newly discovered viral graphs, which were previously unlisted (non-existing), are highlighted in blue.

The 273 structures range in nucleotide length from 13 to 600 monomers. After applying the dual graph representation (see Methods for detailed definitions), corresponding graph IDs were generated and subjected to the dual graph partitioning algorithm (12), resulting in 339 graphs and subgraphs overall. Subgraphs or connected subcomponents of the parent graph were also collected (339 total). Our dual graph partitioning algorithm keeps junctions and pseudoknots intact, as shown in Figure 2. This produced 37 distinct graph entities, including parent graphs and subgraphs, as shown in Figure 4.

**Figure 4:**
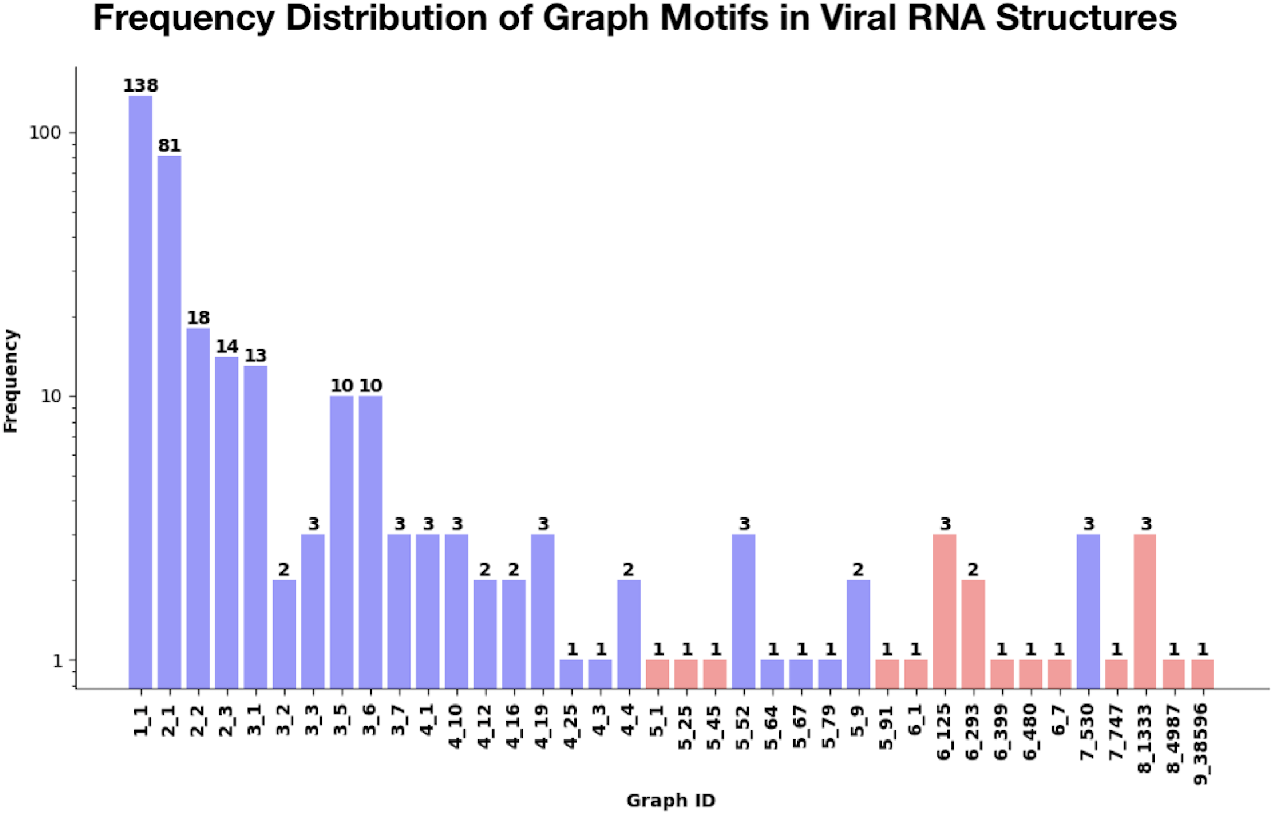
Composition by dual graph motifs of all 339 structures extracted from viral RNAs found in the PDB. The height of each bar indicates the frequency of the corresponding motif in the viral graph library. Newly discovered graphs are highlighted in red. Frequency is described in logarithmic scale.

From Figure 4, we see that the most common structure is 1_1 (138 graphs), followed by 2_1, 2_2, 2_3, and 3_1. Graphs 1_1 and 2_1 represent some of the simplest secondary structures, a stem loop (hairpin) or stem-loop stem (double hairpin) respectively. A hairpin loop consists of a variable length double-stranded RNA (possibly containing mismatched nucleotides) and a terminal apical loop. Hairpin loops appear to have evolved independently in many RNA pathways across a multitude of processes, including mRNA localization, protein binding sites, ribozyme elements, and gene silencing pathways (34; 14). In the 273 viral RNA structures, single or double hairpin loops were present in the trans-activating response (TAR) element of HIV-1 (PDB ID: 6XH2), Hepatitis C IRES (PDB ID: 2KTZ), reverse transcriptase initiation complexes (PDB ID: 6WB2), and many others.

The smallest pseudoknot-containing graph is 2_3 (Figure 2), and was found in various different phylogenetic and functional contexts, such as in an Influenza A virus splicing site at the 3^*′*^ end (PDB ID: 7RQ5), a Luteoviral RNA ribosomal frameshifting element (PDB ID: 2A43), and a Poxvirus RNA in a transcription initiation complex (PDB ID: 7AOZ).

Of the 37 distinct graphs, 14 new motifs emerged with vertex counts ranging from 4 to 8, shown in Figure 5. Of these 14 new graphs, 8 appear as parent graphs, and 6 as subgraphs after partitioning (Graph ID: 4_3, 5_1, 6_125, 7_530, 7_747, 8_4987) (Figure 5). Two new graphs, 6_1 and 6_480, represent multiple non-contiguous RNA strands that serve as a backbone for a Satellite tobacco Mosaic virus capsid protein (33). While the RNA strand of the virus is linked through double helical segments in nature, they were modeled as disjoint RNA strands due to their association with a crystallographic asymmetric unit in the observed experimental data (33). Here we do not distinguish between graphs and subgraphs due to the fact that many PDB structures are variable in length and are truncated from larger RNA structures at varying distances away from functional sites.

**Figure 5:**
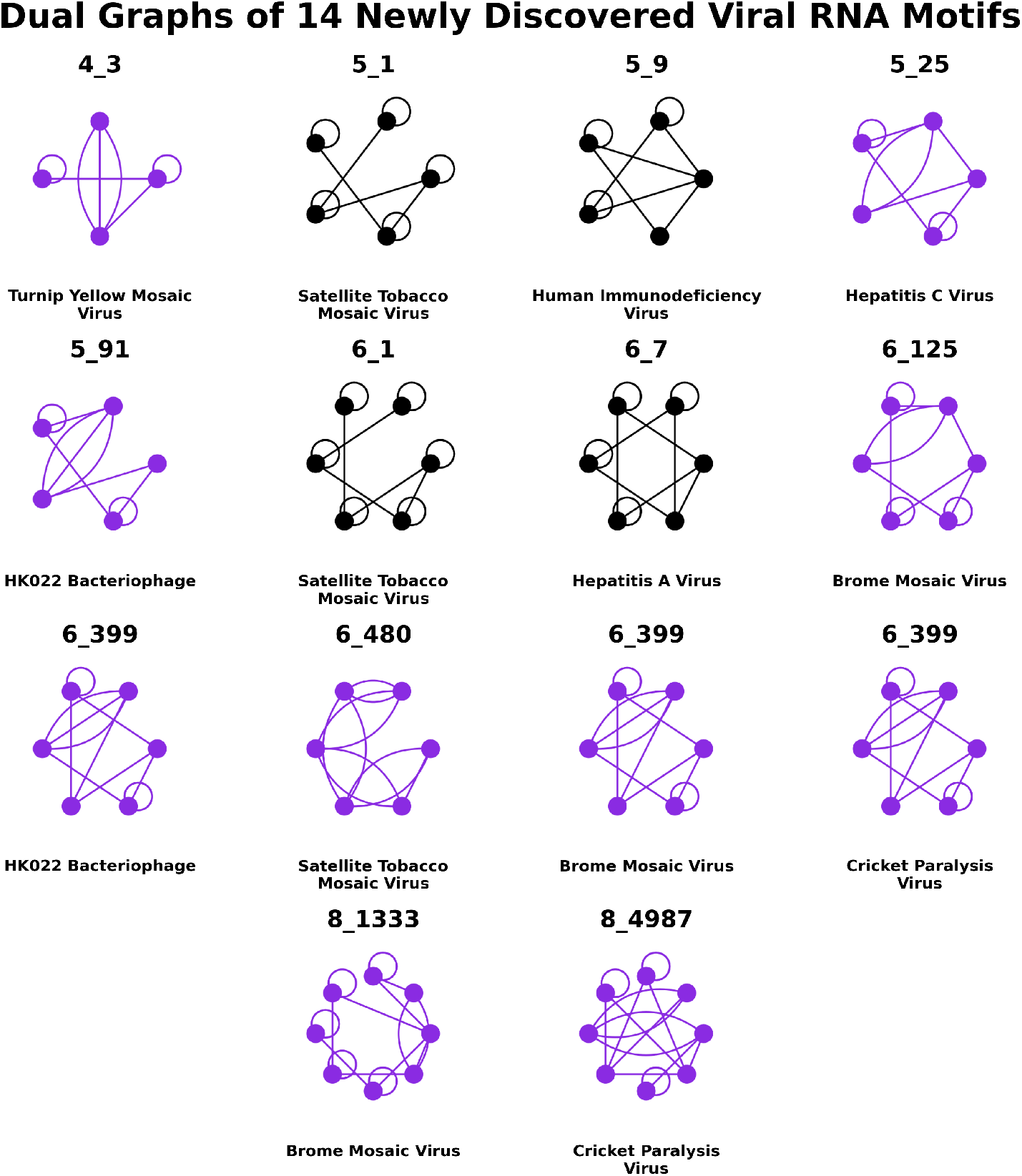
Dual graphs of 14 newly discovered viral motifs, which were not listed in the previous RAG atlas. Graphs containing pseudoknots are highlighted in purple.

**Figure 6:**
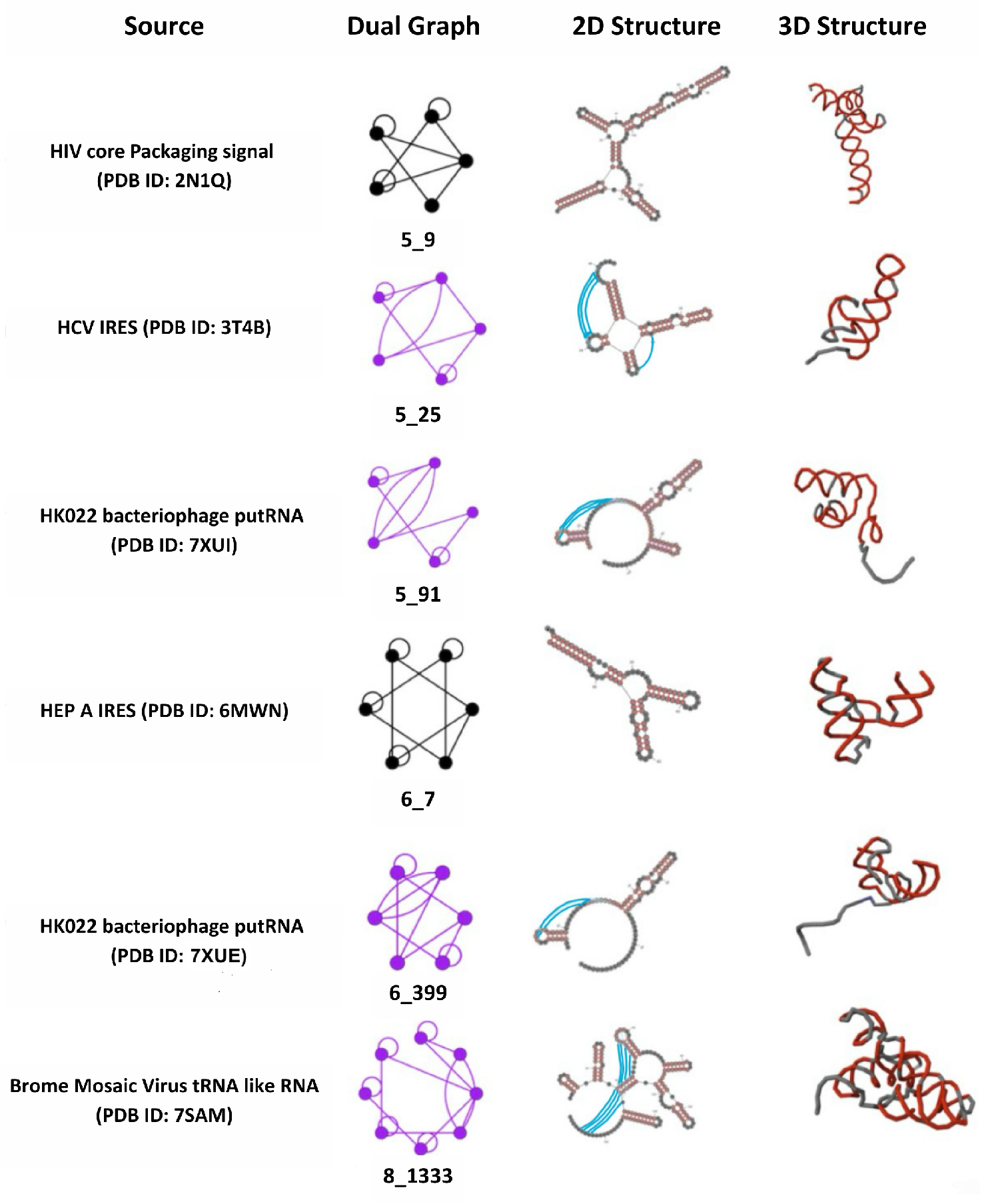
Some new motifs of viral RNA elements from Figure 5 with their dual graphs, 2D, and 3D structures. Viral RNAs analyzed include trans-activation response elements, exonuclease-resistant domains at the 5^*′*^ and 3^*′*^ ends, reverse transcriptase binding domains, RNA polymerase interaction domain riboswitch elements, frameshifting elements, internal ribosome entry sites, core packaging signals, and self-cleaving ribozymes. The two HK022 putRNA examples correspond to distinct related deposited structures from the same study (PDB IDs 7XUE and 7XUI). Red nucleotides indicate base pairs forming stems.

**Figure 7:**
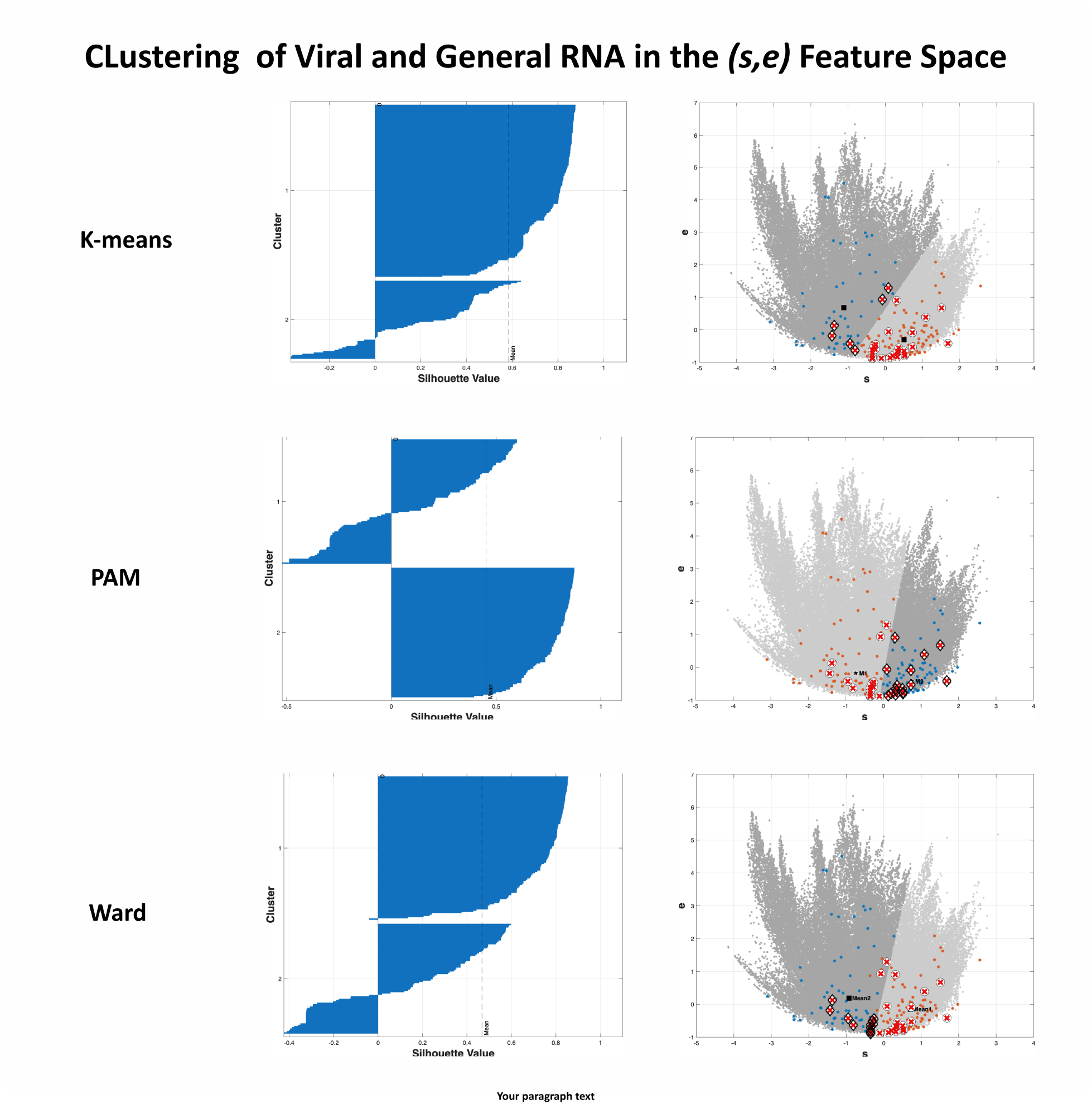
Two cluster partitions of known RNAs in the (*s, e*) feature space (C1 is the cluster with more viral IDs), with corresponding silhouette plots (bottom). Cluster centers are shown as black squares. General RNAs assigned to C1 are red circles and to C2 are blue circles. Viral RNAs assigned to C1 are red crosses and viral RNAs assigned to C2 are black diamonds overlaid with black crosses. Hypothetical RNAs are projected to the nearest cluster centers in light or dark gray. Silhouette plots summarize overall and per–cluster cohesion for each method.

### 2.2 Comparing Viral and General RNA Motifs using Fiedler Score Based Clustering

**Table 1:**
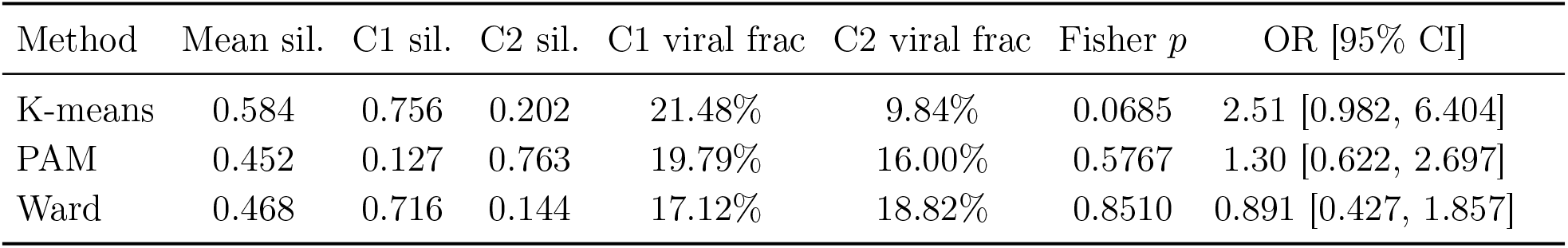
Clustering of known RNAs in the (*s, e*) space. C1 is the cluster containing more viral IDs.

To assess the extent to which viral RNAs share features (submotifs) found in general RNA, we analyzed 196 known RNAs after omitting the single hairpin structure 1_1. Here, *s* (slope) and *e* (residual error) are summary features computed from the ordered, rescaled entries of each graph’s Fiedler eigenvector (see Methods). Of these, 161 are general and 35 are viral, giving a baseline viral RNA prevalence of 17.86% (35/196). We applied three two-cluster algorithms (k-means, k-medoids/PAM, and Ward’s linkage). For each method, we designated the cluster containing more viral IDs as viral (*C*_1_) and the other as general (*C*_2_). Hypothetical motifs were projected to the nearest centroid/mean for visualization only and were excluded from all statistical tests. We use a two-sided Fisher’s exact test to assess whether cluster membership and viral label are associated overall. We also use a right tailed Fisher’s exact test to ask, for each cluster, whether its viral fraction exceeds the baseline viral prevalence of 17.86%. We also report silhouette scores of each cluster, which measures how well separated a cluster is, with values ranging from −1 (misclassified) to +1 (well separated) (28).

For k-means, cluster 1 contained 135 RNAs with 29 viral (21.48%) and had a one-sided Fisher *p* = 0.0346. Cluster 2 contained 61 RNAs with 6 viral (9.84%) and had *p* = 0.988. The global two-sided Fisher test gave *p* = 0.0685 with odds ratio 2.51 and 95% confidence interval [0.982, 6.404]. The mean silhouette was 0.584 (*C*_1_ = 0.756, *C*_2_ = 0.202).

For PAM, cluster 1 contained 96 RNAs with 19 viral (19.79%) and had *p* = 0.306. Cluster 2 contained 100 RNAs with 16 viral (16.00%) and had *p* = 0.810. The global two-sided Fisher test gave *p* = 0.5767 with odds ratio 1.30 and 95% confidence interval [0.622, 2.697]. The mean silhouette was 0.452 (*C*_1_ = 0.127, *C*_2_ = 0.763).

For Ward’s linkage (*K* = 2), cluster 1 contained 111 RNAs with 19 viral (17.12%) and had *p* = 0.692. Cluster 2 contained 85 RNAs with 16 viral (18.82%) and had *p* = 0.450. The global two-sided Fisher test gave *p* = 0.851 with odds ratio 0.891 and 95% confidence interval [0.427, 1.857]. The mean silhouette was 0.468 (*C*_1_ = 0.716, *C*_2_ = 0.144).

Across methods, we find no significant association between cluster assignment and viral label. Taken together, these analyses support a sharing of many common structural building blocks between viral and general RNAs

### 2.3 Shared Motifs Across Viral and General RNAs

Consistent with the lack of separation following motif clustering, we note many similar dual graph motifs used by viral and general RNAs, as shown in Figure 8. In several cases, the same dual graph appears across contexts. For example, graph 4_19 corresponds to cellular tRNA and also appears in viral settings, like the bacteriophage pRNA (PDB ID: 6JXM) and Coxsackie virus replication element (PDB ID: 8DP3).

**Figure 8:**
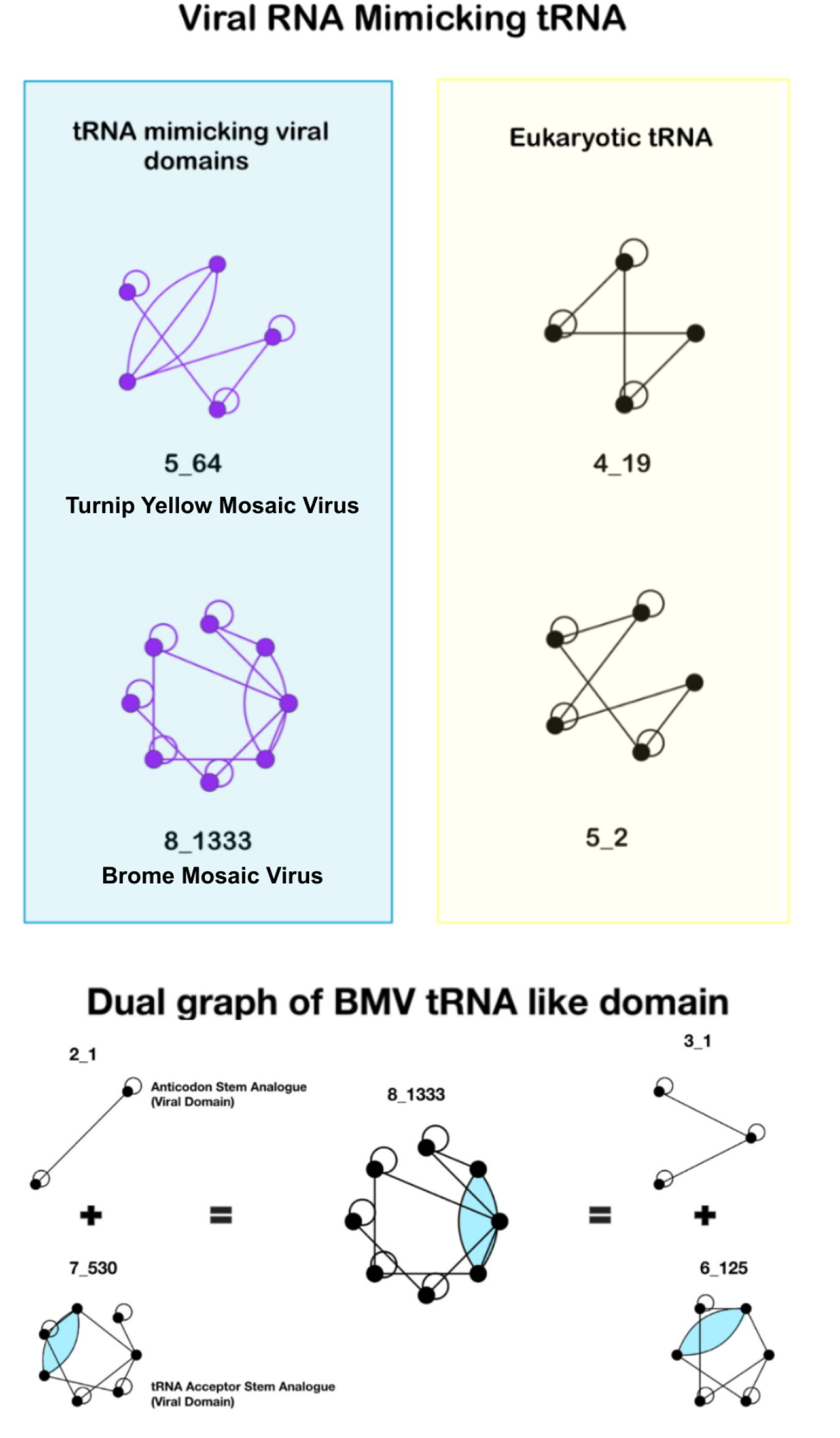
Comparison of viral tRNA mimicking domains and eukaryotic tRNA structures (top) and partitioning of BMV tRNA like domain (bottom). The top left column shows viral tRNA-like motifs, with graph 5_64 representing the tRNA-like structure from Turnip Yellow Mosaic Virus (PDB ID: 4P5J) and graph 8_1333 from the Brome Mosaic Virus tRNA-like domain (PDB ID: 7SAM). The right column displays eukaryotic tRNA motifs, such as 4_19, which was also found to be present in bacteriophage RNA. Graph 5_2 is also included, as it featured in our earlier update as representing a eukaryotic tRNA motif and has not yet been observed among viral graphs.

Specific motif families, like hairpins and double hairpins with IDs 1_1 and 2_1 are prevalent in both sets; viral examples include HIV-1 TAR (PDB ID: 6XH2) and HCV IRES hairpins (PDB ID: 2KTZ). Multi-way junctions show a similar pattern. Viral IRES domains are mapped to dual graphs 5_25 (HCV), 6_7 (HAV), and 9_38596 (CrPV). Together, these examples illustrate that a common translation-initiation role can be realized through distinct dual-graph topologies, including both pseudoknot-free and pseudoknot-containing motifs.

The junction containing subgraph 4_10 occurs in a bacterial riboswitch aptamer and *T. thermophilus* tRNA and also in a viral RNA as subgraph of the HAV IRES graph 6_7.

In addition, graph 6_236 (present in E. coli rRNA) lies near 9_38596 and is the closest non-viral graph to 9_38596 using our scoring model (See Methods). Exoribonuclease resistant folds such as those found in flaviviral viruses like Zika (5_45) and Taman Bat virus (5_67) exhibit junction plus stem layouts, and the same motif recurs in non-viruses, as reflected by 5_45 in the *B. anthracis* ribozyme (PDB ID: 3G9C).

Partitioning further reveals the reuse of motifs across RNAs. The CrPV IRES 9_38596 admits two alternative partitions, 7_747+3_3 and 8_4897+2_3, with 2_3 and 3_3 serving as a common pseudoknot subgraphs in the general atlas (Fig. 2). The BMV tRNA-like domain 8_1333 partitions into 7_530 and the widely used double hairpin 2_1 (Fig. 8). These partitionings of viral RNA show that viruses utilize general RNA submotifs.

Across these examples, variation is mainly in local geometry, such as branch or stem lengths, rather than the underlying dual graph topology. Taken together, these qualitative comparisons align with the clustering result, that viral and general RNAs draw on a shared secondary structure repertoire.

## 3 Summary and Discussion

The development and expansion of the RNA-as-Graphs Motif Atlas, as well as the introduction of the dual graph partitioning algorithm and Fiedler vector derived features, have enabled analysis of the structural intricacies of RNA secondary structure. In this study, using dual-graph representations that preserve pseudoknots and junctions, we described and analyzed existing viral RNA motifs using dual graphs.

We extended the RAG Motif Atlas (Figure 3) by adding 14 novel viral dual graphs (Figure 5) derived from 273 structures, bringing the library to 197 unique topological motifs. These new motifs span internal ribosome entry sites, tRNA-mimicking domains, exoribonuclease-resistant RNAs, frameshifting elements, reverse-transcription initiation elements, and packaging/core signals. The distribution of motifs is skewed toward simple hairpins (1_1, 2_1), with rarer junction and pseudoknot motifs, as shown in Figure 2 for the Cricket Paralysis Virus IRES (PDB: 4D5N).

Our clustering using Fiedler vector features with k-means, PAM, and Ward found a large intersection between viral and general RNA motifs. Thus, viral and general RNAs overlap in this feature space. Correspondingly, qualitative comparisons of recurrent motifs across different contexts, along with dual-graph partitioning, agree with this picture. Some viral structures break into submotifs that are already common in the atlas, and the same secondary structures recur often across viral and non-viral sources.

The atlas extension and clustering results suggest that new viral entries fit naturally within the existing dual graph universe. Motif families such as hairpins, double hairpins, and multi-way junctions are abundant in both viral and cellular contexts, and several viral cases use the exact same graph IDs seen in general RNAs. In some cases where IDs differ, partitioning reveals shared submotifs, and completely new viral graphs may be close in feature space to already existing graphs. This recurrence helps explain why the viral RNAs do not cluster separately.

As more RNA data are analyzed and cataloged as graphs, it is possible that some of the 14 newly identified viral RNA motifs (Figure 5) might also be assigned to general RNA structures. This complexity not only highlights the challenges of predicting RNA function based solely on structural motifs but also opens up new avenues for understanding how RNA diversity is leveraged across biological systems to fulfill a wide array of roles.

Dual graph partitioning highlights modular aspects of RNA structure by decomposing larger topologies into recurrent subgraphs while keeping junctions and pseudoknots intact. In this sense, the resulting subgraphs can be viewed as structural building blocks within larger RNAs. For example, in Figure 8, the 8_1333 graph of the BMV tRNA-mimicking domain partitions into subgraphs 7_530 and 2_1. Here, 7_530 contains the acceptor-stem-like pseudoknotted region, whereas 2_1 corresponds to the double-hairpin portion associated with the putative anticodon-stem analogue. The larger viral motif can be understood in terms of smaller structural units with identifiable roles. More generally, partitioning helps place large viral RNA motifs within a broader repertoire of shared RNA substructures and recurrent modules. Likewise, RNA folding studies have shown that local secondary-structure rearrangements and dynamics shape the behavior of larger RNAs. (39; 24; 45; 46).

Limitations include reliance on deposited structures to the Protein Data Bank with uneven coverage across viral families and the use of two-cluster partitions that were chosen to directly test separability rather than to discover multiple groups. Our analysis also focuses on secondary structure topology. It does not incorporate proteins, ion conditions, or long range tertiary interactions that can shift function within a shared topology.

In addition, graph assignment can depend on the completeness of deposited structures and on secondary-structure annotation, both of which can affect the resulting topology for some RNAs. More generally, dual graphs are flexible coarse-grained representations, and changes in base-pair annotation, minimum stem definition, loop/bulge threshold, or helix-merging rules may change the resulting topology.

In summary, this study contributes to the expanding field of RNA research and RNA structural determination by providing a mathematical analysis of RNA secondary structures as graphs for viral RNA. More generally, identifying recurrent RNA topologies and submotifs may be useful for RNA nanotechnology and for therapeutic efforts that rely on modular RNA structural elements. Examples include antiviral strategies based on targeted RNA mutations, in the same spirit as mutant designs for frameshifting elements of SARS-CoV-2, where confirming experiments were also reported (19; 20; 43), HIV-1 (47), and Chikungunya (48).

## 4 Methods

### Dual Graph Definitions

Dual graphs were defined by representing RNA double-stranded stems as vertices and intervening single-stranded regions as edges. A stem containing at least two consecutive base pairs is represented as a vertex, and isolated single base pairs are ignored. Hairpin loops are represented as self-edges. Junctions connecting two helices are represented as edges between the corresponding vertices. The dangling 5^*′*^ and 3^*′*^ ends are ignored. Small interruptions between adjacent base pairs are handled during graph construction as described below. In this implementation, neighboring helices separated by bulges or internal loops of up to 5 nucleotides are merged into a single stem vertex, whereas larger interruptions are treated as separating adjacent helical stems into distinct vertices connected by the corresponding loop or junction edge. (11; 24)

### RAG Library

The RAG library can be accessed through http://www.biomath.nyu.edu/?q=rag/home. By iteratively connecting smaller graphs, 110,668 dual graphs of 1-9 vertices have been enumerated (15). The RAG library organizes dual graphs by vertex count *V*, which are then sorted by ascending order of their Fiedler values. Each graph is assigned an ID *V*_*n*_, ordered for each *V* by increasing compactness.

### RAG Framework

RAG defines 2D RNA structures as tree or dual graphs. The RAG framework includes tools for subgraph partitioning (12), and clustering and scoring models (15; 13). Partitioning is valuable because of the inherent modularity of RNA structures (13). The RAG partitioning decomposes larger parent dual graphs into connected subgraphs, or irreducible graph modules, using spectral graph properties, including the Fiedler-vector framework. Additionally, partitioning graphs can enhance the efficiency of computational algorithms for RNA structure prediction and comparison, aiding graph partitioning in the systematic exploration of RNA architecture and its biological implications. The graph partitioning was improved in 2019 (14) with the introduction of an algorithm that preserves pseudoknots and junctions, as illustrated by partitioning one large virus (Figure 2). The resulting modules are interpreted as recurrent RNA submotifs within larger RNA architectures.

The dual graph enumeration algorithm allows us to expand the RAG atlas to 2288 tree graphs of 1–13 vertices and 110,668 dual graphs of 1–9 vertices (15). Graphs which have real corresponding (experimentally resolved) RNA structures are denoted as ‘existing’. Here we refer to the 183 previously existing RNA as general RNAs, to distinguish them from the viral RNA motifs we explore. Here, graphs which have yet to be assigned a real RNA structure are called ‘hypothetical’.

### Graph Representation

Secondary structures in dot-bracket representation were extracted from PDB structures using 3DNA-DSSR (40) and converted into connectivity table (CT) files using `dot2ct` from the RNAstructure package (41). CT files were processed using `dualGraphs.py`, based on the Schlick laboratory’s `Existing-Dual-Search` repository. The script extracts CT base-pairing patterns, groups consecutive paired residues into helices, assigns helices as vertices, and assigns single-stranded regions connecting helices as edges. For this analysis, we used a lightly modified version of the helix-merging step in `dualGraphs.py`, merging helices separated by interruptions of up to 5 nucleotides. Isolated single-base-pair helices are removed. The resulting adjacency matrix is matched with the RAG dual-graph library to assign the corresponding graph ID (11; 24). Because the dual graph is constructed from the annotated base-pairing pattern, alternative base-pairing annotation methods could alter the inferred helices, loops, and junctions, and therefore change the assigned dual graph for some RNAs.

In order to extract dual subgraphs from newly assigned structures, we used `subgraphs.py`. Starting from an assigned graph ID, `subgraphs.py` retrieves the corresponding adjacency matrix, partitions the graph into irreducible components (see below under Dual Subgraphs/ Partitioning), and assigns graph IDs to the resulting subgraphs by matching them back to the RAG dual-graph library.

### Laplacian and Fiedler Eigenvector

Every graph with *n* vertices can be described by an *n* × *n* adjacency matrix *A* and entries *a*_*ij*_, with *a*_*ij*_ equal to the number of edges between vertices *i* and *j*, and *a*_*ii*_ equal to 2 for self-connected vertices. There also exists for every graph a degree matrix *D* where the *a*_*ii*_ entry is equal to the number of incident edges on vertex *i*. The Laplacian *n* × *n* matrix *L* = *D* − *A* is calculated by taking the difference between a graph degree matrix and its adjacency matrix. The Laplacian is a positive semidefinite matrix and as such has non-negative eigenvalues with *λ*_1_ = 0 being the smallest eigenvalue with corresponding eigenvector *µ*_1_ = (1, 1, …, 1). The focus is on the second smallest eigenvalue *λ*_2_, also known as the Fiedler value, which represents the algebraic connectivity of the graph, and its corresponding eigenvector, the Fiedler vector (42). A higher Fiedler value suggests greater algebraic connectivity.

### Dual Subgraphs/Partitioning

An articulation point is a vertex whose removal, along with the associated edges, results in a disconnected graph. The dual graph partitioning algorithm preserves connected components by only partitioning graphs along articulation points. The partitioning algorithm leverages the principles of a gap cut algorithm, which efficiently partitions graphs into weakly inter-connected regions, which are joined by an articulation point (44). In order to partition a graph, its Fiedler vector is derived and the gap cut algorithm identifies the largest ‘gap’ between the Fiedler vector components, which correspond to vertices of the graph. This also preserves pseudoknots, junctions, and internal loops which are often closely related to RNA function (12). A subgraph is the resultant graph after partitioning, and can be thought of as a connected component, or a recurrent structural submotif, within the larger parent RNA graph. An example is given in Figure 1.

### Fiedler Scoring and Clustering

Although the Fiedler value provides insight into the graph’s algebraic connectivity, which indicates how tightly the graph is connected, it is not always sufficient for distinguishing between graphs that have similar connectivity but differ in their structural composition. We extract additional features by analyzing the entries of the Fiedler vectors that help to capture more information about the graph’s structure. For a graph of *n* vertices, there is a one-to-one correspondence between vertices and the entries *µ*_2,*i*_ of the Fiedler vector (*µ*_2,1_, *µ*_2,2_, …, *µ*_2,*n*_), which is unique for each Laplacian up to change of sign. The arrangement of vertices in a graph is reflected in the distribution of the Fiedler vector components *µ*_2,*i*_. Common topological modules in dual graphs show distinct patterns in this distribution. In particular, vertices in linear modules tend to have Fiedler vector components that increase monotonically (13). Similarly, vertices forming *K*-way junctions with similar free-end branches often share the same Fiedler vector component values. When taken as an ordered set, these components tend to increase monotonically in common RNA dual graph modules. Therefore, given that adjacent vertices have close *µ*_2,*i*_ values, and these values increase from one end of the graph to the other, analyzing the distribution of the ordered components provides additional structural insights, which are then used to cluster graphs (13).

### Features *s* and *e*

The scoring and motif search algorithms follow the approach outlined in the 2021 paper (13). The clustering process itself relies on parameters *s* and *e* which are derived from the sorted and rescaled entries of each graph’s Fiedler vector *µ*_2_ (second eigenvector of the Laplacian associated with the graph). To calculate these parameters for a particular graph:

1. Calculate the normalized Fiedler vector *µ*_2_ and arrange the components *µ*_2,*i*_ in ascending order.
2. Scale each *µ*_*i*_ by:

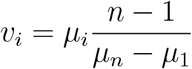
3. Perform linear least squares regression on the points (1, *v*_1_), (2, *v*_2_), …, (*n, v*_*n*_) to obtain the slope *s* and the mean square error *e*. For each graph *i*, the planar coordinates (*s, e*) are plotted.

### Scoring Model

The scoring model is constructed by incorporating the extracted features *s* and *e*, which are plotted on a plane, along with the graphs’ associated weights from a training set which defines the clustering.

An ‘initial score’ for a graph *i* in the training set, *ES*_*i*_, is calculated based on the weight *w*_*i*_ of the graph *i*, and *σ* and *ϵ* are freely chosen parameters to control the influence of the weight on the score:

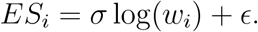

Each graph *i*, represented as a point in the plane, contributes to the score of a graph *j* based on their Euclidean distance *d*_*ij*_, via the pairwise score *S*_*j,i*_:

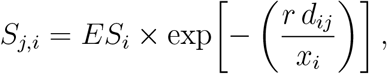

where *r* controls the effect of weight on the pairwise score and *x*_*i*_ is a scale parameter.

For each graph *j*, the total score it receives is

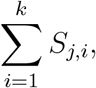

where *k* is the total number of graphs in the training set.

### Clustering

We clustered (*s, e*) points with three standard methods: *K*-means (25), PAM (26) and Ward’s linkage (27). For all methods we set *K* = 2; for Ward we cut the dendrogram at *K* = 2. Clusters are summarized by the mean silhouette overall and per cluster (28). Specifically, *K*-means solves

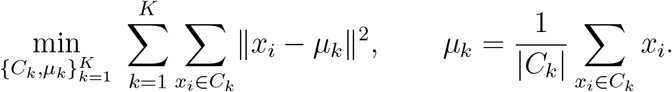

PAM selects medoids *m*_*k*_ ∈ *C*_*k*_ to minimize

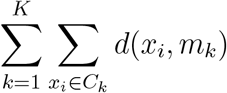

for a chosen dissimilarity *d*(*x*_*i*_, *x*_*j*_) = ∥*x*_*i*_ − *x*_*j*_∥_2_.

After fitting each method, we relabel clusters so that Cluster 1 (C1) is the cluster containing the larger number of viral IDs among the known RNAs, and Cluster 2 (C2) is the other cluster. We report the overall mean silhouette and the per-cluster mean silhouettes for C1 and C2 (28).

Clustering was performed in the (*s, e*) feature space, where *s* and *e* are derived from the sorted and scaled components of the graph Fiedler vector. Together, these features summarize the distribution of Fiedler-vector entries and thus encode coarse-grained properties of the assigned dual-graph topology. (13)

We did not assign separate weights to different structural features after graph construction. All features were analyzed only through the final dual-graph topology and the associated Fiedler-based descriptors.

### Significance testing

To assess whether cluster membership and viral label are associated among known RNAs, we applied Fisher’s exact test on the 2 × 2 table of label (viral, general) by cluster (C1, C2). For each cluster separately, we additionally tested whether its viral fraction exceeded the overall viral fraction using a right-tailed Fisher exact test. We report *p*-values, odds ratios, and 95% confidence intervals.

## Available Code

The dual-graph construction workflow used in this work was based on the Schlick laboratory’s `Existing-Dual-Search` repository, available at https://github.com/Schlicklab/Existing-Dual-Search. The general dual-graph construction code is provided in the repository as `dualGraphs.py`. For this work, we used a slightly modified version provided in `Existing-Dual-Search/Jad/dualGraphs_ILmax5.py`, which merges neighboring helices separated by interruptions of up to 5 nucleotides. Graph partitioning and subgraph identification were performed using `subgraphs.py`. Clustering analyses were performed in MATLAB using standard implementations of *K*-means, PAM, and Ward’s linkage.

## 5 Acknowledgments

We gratefully acknowledge funding from the National Science Foundation Award DMS-2151777 and CHE-2330628 from the Division of Mathematical Sciences and the Division of Chemistry, the National Institutes of Health R35GM122562 Award from the National Institute of General Medical Sciences, and the Philip Morris International. Support from the Simons Foundation through the NYU Simons Center for Computational Physical Chemistry Award MPS-T-MPS-00839534 to Jad Zbib and Tamar Schlick is also gratefully acknowledged.

